# A Comprehensive Approach Characterizing Fusion Proteins and Their Interactions Using Biomedical Literature

**DOI:** 10.1101/371088

**Authors:** Somnath Tagore, Alessandro Gorohovski, Lars Juhl Jensen, Milana Frenkel-Morgenstern

## Abstract

Today’s increase in scientific literature requires the efficient methods of data mining for improving the extraction of the useful information from texts. In this manuscript, we used a data and text mining method to identify fusions and their protein-protein interactions from published biomedical text. The extracted fusion proteins and their protein-protein interactions are used as a training set for a Naïve Bayes classifier that is further used for final identification of testing dataset, consisting of 1817 fusions. Our method has a literature corpus, text and annotation mappers; keywords, rule bases, negative tokens, and pattern extractor; synonym tagger, normalization, regular expression mapper; and Naïve Bayes classifier. We classified 1817 unique fusion proteins and their corresponding 2908 protein-protein interactions for 18 cancer types. Therefore, it can be used for screening literature for identifying mentions unique cases of fusions that can be further used for downstream analysis. It is available at http://protfus.md.biu.ac.il/.

## 1 Introduction

### 1.1 Background

Fusion proteins resulting from chromosomal translocations play important roles in many types of cancer and are extensively discussed in the cancer literature. However, because they do not exist as entities in the normal, non-cancer genome, they are usually not considered when text mining the biomedical literature.

To collect information about fusions, we have an in-house database, ChiTaRS [1] that covers more than 11,000 fusion breakpoints. However, this resource is still incomplete and needs to kept up-to-date with the ever-growing literature. We thus set out to extract the unique fusion proteins associated with different cancer subtypes along with their protein–protein interactions from PubMed abstracts.

Currently, PubMed comprises more than 28 million citations, with approximately 14000 cancer-related papers from 2018 alone and more than 3 million abstracts in total that mention cancer. However, finding the fusion proteins mentioned within these is non-trivial, because a fusion protein such as BCR–ABL1 can be represented in variable forms in the text [2]. These variations include the formatting of the fusion instances themselves (e.g., BCR-ABL1 vs. BCR:ABL1 vs. BCR/ABL1 vs. BCR-ABL1), and the keywords used to describe that they are fusions (fusions vs. fusion proteins vs. chimers vs. chimeras) [3]. Moreover, when extracting protein–protein interactions of fusions, their actions can be described in varying ways (e.g., activate vs. interact vs. express vs. induce [4]. ***Fig 1*** provides the overall methodology of ProtFus.

**Fig 1:**
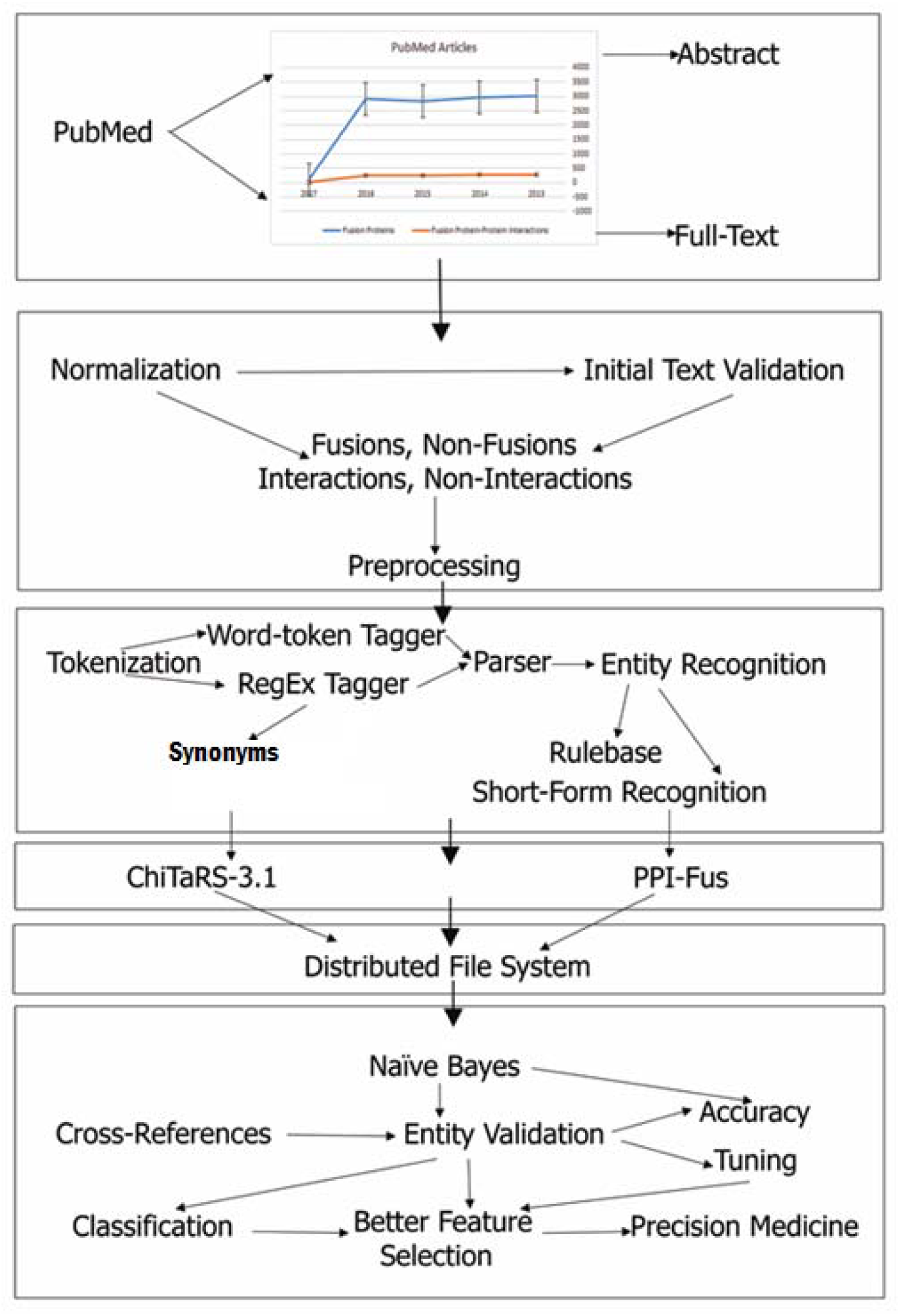
The overall methodology of ProtFus. The algorithm begins with collecting abstracts and full-text from PubMed; followed by normalization; tokenization, entity recognition; cross-references; databases; and machine learning classifier.

### 1.2 Previous studies in the field

Some previous work has been carried out on developing text mining approaches for modern medical text, especially cancer. For example, several annotated corpora have been created [5-7]. Likewise, various supervised-based methods have been developed, using linguistic patterns for detecting semantics in gold standard corpora, e.g., part-of-speech (nouns or verbs). NEs frequently consist of such sequences of nouns and adjectives, while NEs involved in relationships use verbs. Further, several tools are now available offering to extract specific information from literature. For example, some well-known biomedical text mining tools include MetaMap [8], WhatIzIt [9], iHOP [10], Gimli [11]. Moreover, continuous evaluations and verifications are done by the biomedical text mining community through BioCreative [12-13], BioNLP [14], i2b2 [15], to name a few. Also, a number of efforts for creating resources accounting for evolving ways have been provided, that are referenced from time-to-time. For instance, for identifying relationships among NEs, there exist certain difficulty in extracting syntactic parses. Such problems can be dealt by means of identifying co-occurrences between the NEs.

Availability of resources having complete information related to fusions in cancers as well as their protein-protein interactions is less [16-17]. There are some well-known databases of fusion proteins, such as ChiTaRS-3.1 [1], ChimerDB 3.0 [18], COSMIC [19] and TICdb [20]. All these databases have limited number of fusions, which are not unique. Further, there are currently no methods that can successfully identify fusions and their interactions from scientific literature. Thus, the success of ProtFus lies in removing these lacunae and predicting unique fusions from the literature. In this scenario, efficient methods of text mining provide us unique ways for the extraction and interpretation of information present in these scientific public resources as follows:

1. We are interested in identifying fusion proteins and their interactions from published scientific articles [21-22].
2. From the point-of-view of text mining, this task deals with identifying information that need to be tagged ‘concurrently’ to find ‘co-mentions’, like human fusion proteins and cancer. Let us assume that we are interested in the fusion protein BCR-ABL1. We want to find all the mentions of BCR-ABL1 in the literature. But, BCR-ABL1 can be spelled in a variety of ways BCR-ABL1, BCR/ABL1, bcr-abl1, bcr/abl1, bcr:abl1, BCR:ABL1, etc. Thus, we need a good ‘tagger’ which can identify all these jargons.
3. Further, we aim to identify interaction occurrences among fusion is trickier as it requires tagging interaction tokens from literature as well as linking them to their corresponding proteins. For instance, in the text ‘Grb2 has been shown to bind NPM-ALK and ATIC-ALK in previous works’, the interaction token is ‘bind’.
4. We have developed Protein Fusions Server (ProtFus), a resource to identify instances of fusion proteins and their interactions from literature [23, 12], based on text mining approaches using natural language processing (NLP) methods [24-25].
5. The major goal is to identify the co-occurrences of both fusions and their corresponding interactions by filtering out the false positives cases from general searches on fusion proteins using PubMed, so that a more focused and restricted result could be generated.

The resulting instances could be further used for the designing methods in Precision Medicine for fusions as cancer drug targets, combining classical natural language processing and machine learning techniques.

## 2 Methods

The basic framework of ProtFus is provided in (***Fig 1***). It consists of the basic computational methods, like text mining, machine learning, and a distributed database system for storing the text as well as features extracted from biomedical literature. Here, we discuss the development of ProtFus to extract fusion information (e.g., BCR-ABL1), protein-protein information (e.g., BCR-ABL1 causes leukemia), prediction models (e.g., Naïve Bayes) for text classification extracted from PubMed references; as well as a list of the cancer fusions from Mitelman Database of Chromosome Aberrations and Gene Fusions in Cancer [26]; and the Breakpoints Collection of the ChiTaRS-3.1 database [1], respectively (***Table 1***).

**Table 1.**
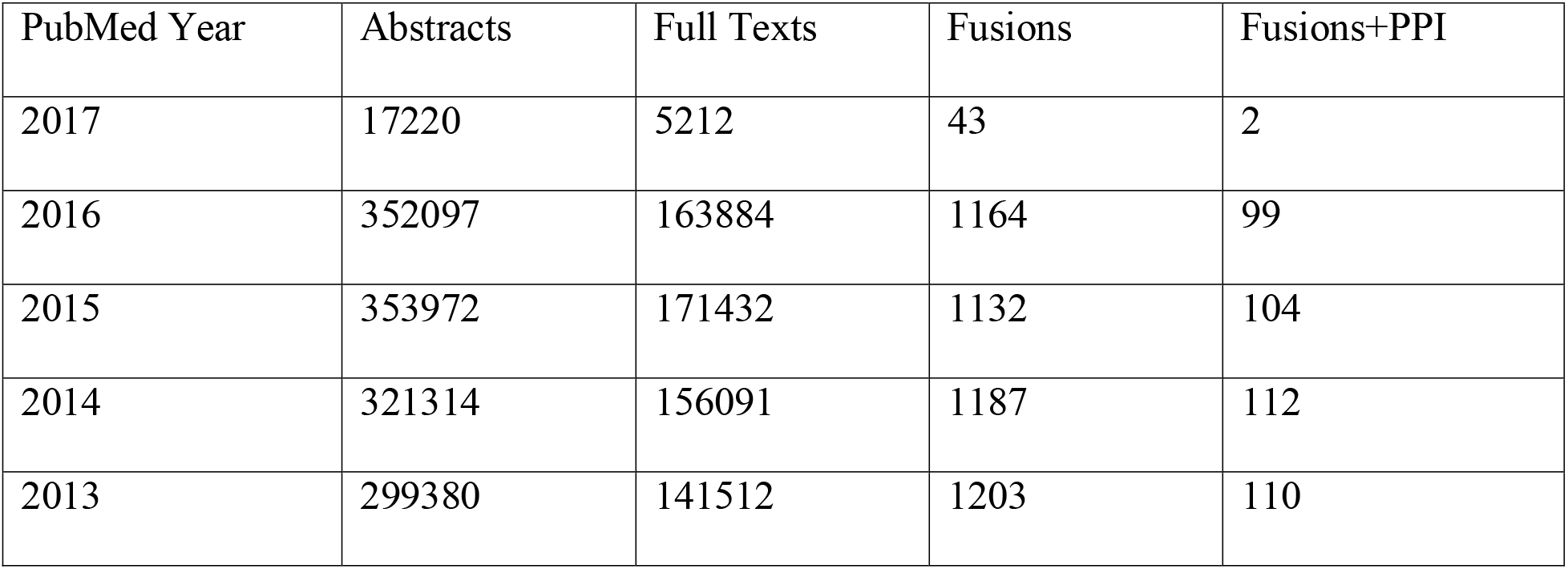
Datasets considered for Training.

| PubMed Year | Abstracts | Full Texts | Fusions | Fusions+PPI |
| --- | --- | --- | --- | --- |
| 2017 | 17220 | 5212 | 43 | 2 |
| 2016 | 352097 | 163884 | 1164 | 99 |
| 2015 | 353972 | 171432 | 1132 | 104 |
| 2014 | 321314 | 156091 | 1187 | 112 |
| 2013 | 299380 | 141512 | 1203 | 110 |

**Table 2.** Datasets considered for Testing ProtFus.

| PubMed Year | Abstracts | Full Texts | Fusions | Fusions+PPI |
| --- | --- | --- | --- | --- |
| 2017 | 25830 | 7819 | 65 | 5 |
| 2016 | 528146 | 245826 | 1747 | 148 |
| 2015 | 530960 | 257148 | 1697 | 155 |
| 2014 | 481971 | 234136 | 1780 | 167 |
| 2013 | 449069 | 212268 | 1805 | 165 |

### 2.1 Initial text validation

The initial text validation is performed for input from PubMed to remove false positive results, followed by segregation into ‘tokens’. We performed stemming of the words for sentences, followed by identifying named-entities within sentences with ‘porter2’ algorithm using ‘stemming’ package in python [27]. The named-entities within sentences were blanked out to make it more generalized. This was followed by using a bag-of-words representation [28] based on a Frequency score (*FS*) for estimating the importance of selecting a token. For the bag-of-words representation, we use the *FS* threshold (*T_s,a_*) Eq (1):

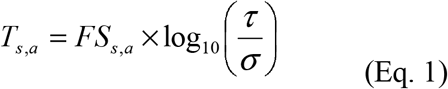

Here, *FS_s,a_* denotes the frequency of token *s* in article *a*, *τ* is the number of articles/abstracts, *σ* is the number of articles having *s*. We used the Naïve Bayes classification method for building a prediction model [28] for categorizing the tokens in abstracts/articles to either fusion proteins or interactions in fusions, as per the Medical Subject Headings (MeSH) items. The ProtFus framework was developed on an Apache cluster, with a back-end My-SQL database and the support from ChiTaRS-3.1 [1]. The tool was developed using Python, whereas the interface was developed using CGI-Perl.

### 2.2 Feature extraction

We use the N-gram model for detecting N-words at a time from a given sentence. An N-gram model model’s sequences, notably natural languages, using the statistical properties of N-grams. This idea can be traced to an experiment by Claude Shannon’s work in information theory [29]. Particularly, using a 2-gram method, all words in a sentence are broken down into two different combinations including unigram and bigrams, i.e., one and two words [30]. For example, some possible sets of combinations are provided in (***Fig 2***). We extracted a set of bigrams as well as combinations of 3-grams and 4-grams from abstracts or full-text articles for training ProtFus to detect specific fusion proteins instances. Along with this, the instances of these tokens were also counted in the back-end corpus [31]. Further, when *FS* was the standard feature score, a considerable high threshold (*T_s,a_*) was given to tokens that appeared frequently in the corpus. Moreover, we also converted all abstracts or full-text articles into ‘similar-length’ feature vectors, where each feature represents a (*T_s,a_*) of the found token.

**Fig 2:**
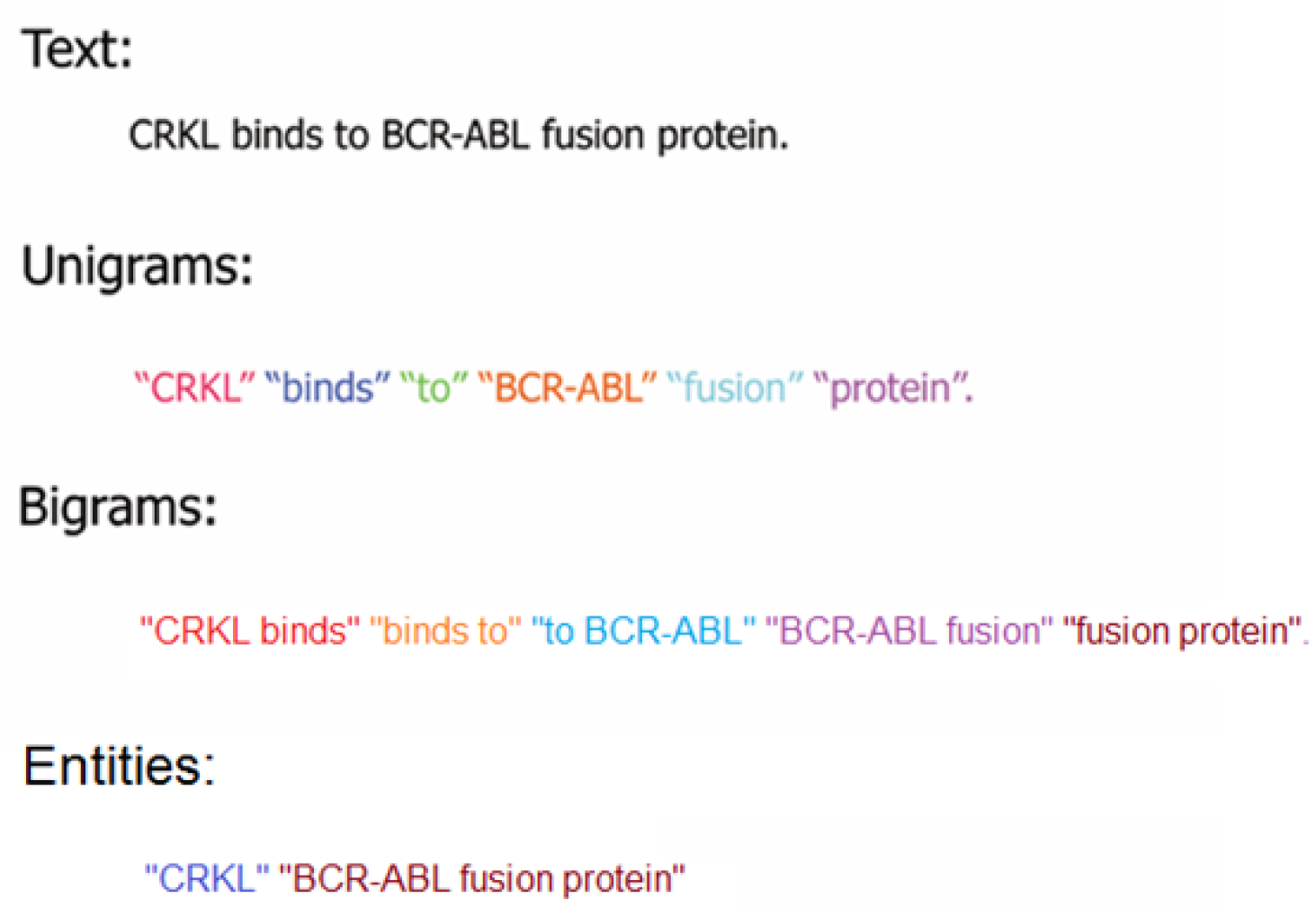
N-gram model for detecting N-words by ProtFus. The N-gram model and some possible set of combinations.

This was followed by organizing a bag-of-words representation of the feature vectors (***Table 3***). Thus, ***Tables S1-S2 (Suppl. data)*** include the back-end corpus considered for tagging fusions and their interactions. The word-token tagger has a back-end Synonyms (with synonyms resource, ***Table S3, Suppl. data***) whereas the RegEx tagger has a back-end Synonyms (with rulebase, ***Table S4, Suppl. data***). Likewise, ***Table 4*** represents Precision and Recall for retrieval step.

**Table 3.** Bag-of-words collection for 10 PubMed ID Abstracts.

| PMID | Fusions | Fusion Gene | Biological Token | Miscellaneous Token |
| --- | --- | --- | --- | --- |
| 24186139 | 1 | 1 | 20 | 35 |
| 22101766 | 0 | 1 | 25 | 30 |
| 18451133 | 0 | 1 | 28 | 38 |
| 11930009 | 1 | 1 | 26 | 32 |
| 15735689 | 0 | 1 | 21 | 34 |
| 18850010 | 0 | 0 | 27 | 33 |
| 21193423 | 1 | 0 | 23 | 33 |
| 22570737 | 1 | 0 | 30 | 38 |
| 18383210 | 1 | 0 | 29 | 35 |
| 24345920 | 1 | 0 | 26 | 32 |
| 16502585 | 1 | 0 | 21 | 33 |

**Table 4.** Precision and Recall for retrieval step.

| Dataset | Precision | Recall | F-Score | Accuracy |
| --- | --- | --- | --- | --- |
| Set A | 0.79 | 0.82 | 0.76 | 0.81 |
| Set B | 0.81 | 0.83 | 0.78 | 0.8 |
| Set C | 0.85 | 0.84 | 0.82 | 0.85 |
| Set D | 0.72 | 0.76 | 0.72 | 0.74 |
| Set E | 0.8 | 0.82 | 0.78 | 0.82 |
| Set F | 0.81 | 0.81 | 0.78 | 0.82 |
| Set G | 0.78 | 0.83 | 0.81 | 0.83 |
| Set H | 0.75 | 0.81 | 0.78 | 0.8 |
| Set I | 0.85 | 0.82 | 0.81 | 0.83 |
| Set J | 0.73 | 0.78 | 0.76 | 0.75 |

### 2.3 Named-Entity Recognition

The tokens have been then used to parse the texts for performing named-entity recognition. Named-entity recognition (NER) locates and classifies named entities in text into pre-defined categories. For example, the unannotated block of text ‘CRKL binds to BCR-ABL fusion protein’ can be annotated as ‘[CRKL]_protein_ binds to [BCR]_protein_ -[ABL]_protein_ [fusion protein]_key_’. This is followed searching for a pattern like, protein1-protein2 key or protein1/protein2 key or protein1:protein2 key. We also perform named-entity recognition of diseases to understand how a fusion may be related to a certain cancer. For example, in ‘BCR-ABL causes leukemia’, we perform annotations as ‘[BCR]_protein_-[ABL]_protein_ [causes]_action-verb_ [leukemia]_cancer_’. A search can be performed by using a PubMed abstract or uploading a text file or based on a specific input text. For example, in case of an input text, the result is displayed in a separate window, with the fusion proteins being highlighted. Similarly, in case of protein-protein interactions among fusions, the result window includes the input text with interactions being highlighted. Another feature of ProtFus includes directly searching using PubMed articles [27]. Users can select from the given drop-down box, the number of articles that need to be considered for searching fusions and their interactions. The result includes the abstracts of all those articles, which match best with fusion proteins keywords. This file can be further used for highlighting the fusions and their interactions. ***Table 5*** represents Precision and Recall for named-entity recognition.

**Table 5.** Precision and Recall for named-entity recognition.

| Dataset | Precision | Recall | F-Score | Accuracy |
| --- | --- | --- | --- | --- |
| Set A | 0.79 | 0.82 | 0.77 | 0.81 |
| Set B | 0.77 | 0.82 | 0.8 | 0.82 |
| Set C | 0.87 | 0.83 | 0.82 | 0.89 |
| Set D | 0.8 | 0.81 | 0.76 | 0.78 |
| Set E | 0.84 | 0.83 | 0.82 | 0.82 |
| Set F | 0.81 | 0.84 | 0.83 | 0.83 |
| Set G | 0.81 | 0.89 | 0.85 | 0.84 |
| Set H | 0.82 | 0.82 | 0.84 | 0.8 |
| Set I | 0.82 | 0.84 | 0.83 | 0.87 |
| Set J | 0.78 | 0.8 | 0.77 | 0.79 |

## 3 Results

The fusion protein data were validated using ChiTaRS-3.1 [1] database of concrete known cases of fusions and interactors as well as Mitelman database of Chromosome Aberrations and Gene Fusions in Cancer. The fusion protein validation was performed by searching the corresponding occurrences of breakpoints in cancers from the ChiTaRS database and comparing them with that of the result provided using ProtFus. The Mitelman database was used mainly for identifying potential fusions, their role in cancer. Similarly, the interactions were validated using the Chimeric Protein-Protein Interaction Server (ChiPPI) [35], STRING database [32] and iHOP database [10, 3]. The interactions information that we got was compared with that of ChiPPI [35] and STRING [32], by performing a simultaneous search in both. Since, we were particularly interested in searching for instances of interactions from scientific literature, we also relied on the iHOP database.

### 3.1 Designing the model for training and testing

We downloaded abstracts and full-text articles from PubMed to generate both training and testing datasets. Separating a dataset into ‘training data’ and ‘testing data’ is an important part of evaluating text classification models. In such a dataset, a training set is used to build up a prediction model, while a testing set is used to evaluate the model built. Training data consists of abstracts as well as full-texts from PubMed database for fusions and their protein-protein interactions. To this end, for each dataset illustrated in ***Table 1***, the 10-fold cross validation, each time using 40% of the entities to train a prediction model and the remaining 60% to test it. We utilized 64-bit Linux CentOS operating system on a cluster platform built with four Dell servers of the 32 compute nodes and configured with 1TB RAM memory each, and 1PetaByte storage.

### 3.2 Tokenization of bag-of-words reveals biological, miscellaneous, function and literals tokens

Tokenization is performed using two specific taggers:

1. Word-token tagger
2. RegEx tagger

The word-token tagger identifies property words from the text for fusions, like ‘fusion proteins’, ‘fusion transcripts’, ‘chimeric proteins’, ‘chimeric genes’, ‘fusion gene transcripts’, etc, and action words for protein-protein interactions (PPIs), like ‘activate’, ‘block’, ‘depend’, ‘express’, ‘interaction’, etc. Similarly, the RegEx tagger recognizes associates these word-tokens with their corresponding literals. For example, given the following text, The small molecule BCR-ABL-selective kinase inhibitor imatinib is the single most effective medical therapy for the treatment of chronic myeloid leukemia (CML), the tokenization output is Biological Tokens: ‘small’, ‘BCR-ABL-selective’, ‘single’, ‘medical’, ‘chronic’; Miscellaneous Tokens: ‘molecule’, ‘kinase’, ‘imatinib’, ‘therapy’, ‘treatment’, ‘myeloid’, ‘leukemia’; Function Tokens: ‘effective’, ‘inhibitor’; Literals Tokens: ‘is’, ‘the’, ‘for’, ‘of’.

The tokenizer module segregates the text into ‘Biological’, ‘Miscellaneous’, ‘Function’ and ‘Literals’ tokens. The biological tokens correspond to those words which are mainly nouns; the miscellaneous tokens correspond to verbs, pro-verbs, adverbs etc; Function tokens correspond to action words or adjectives; and Literals correspond to conjunctions. The process of tokenization is a very important step in our script, as it filters out essential tokens (like protein and function tokens) from non-essential ones (like miscellaneous and literals) for better extraction. ***Tables 4-5*** represent the Precision and Recall for retrieval step and named-entity recognition, respectively. Similarly, ***Table 6*** provides the overall accuracy of the Naïve Bayes classifier, whereas ***Table 7*** represents a comparative analysis of overall prediction rate of fusions and their PPI among ProtFus and some other resources.

**Table 6.**
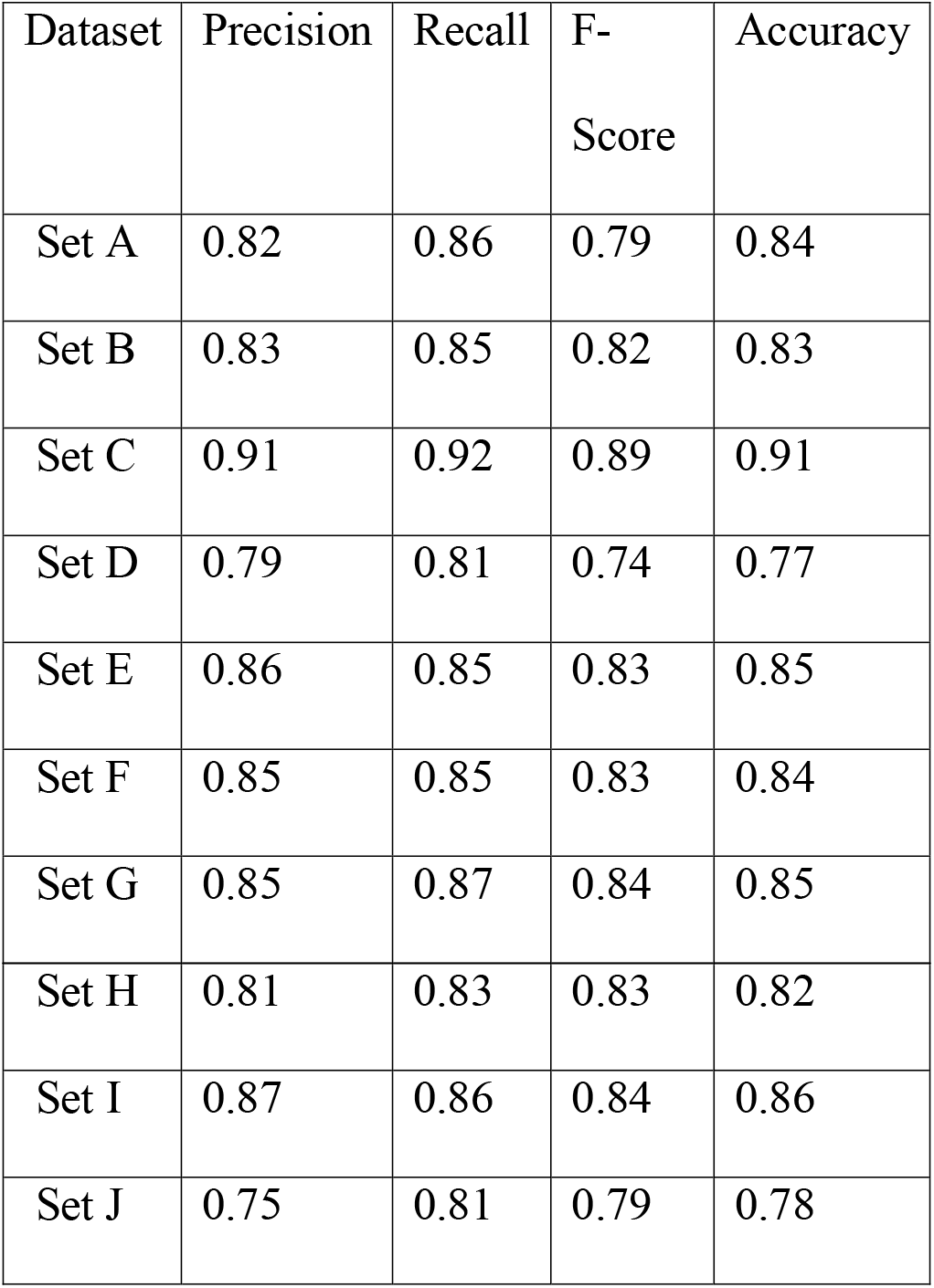
Accuracy Score of Classifier.

| Dataset | Precision | Recall | F-Score | Accuracy |
| --- | --- | --- | --- | --- |
| Set A | 0.82 | 0.86 | 0.79 | 0.84 |
| Set B | 0.83 | 0.85 | 0.82 | 0.83 |
| Set C | 0.91 | 0.92 | 0.89 | 0.91 |
| Set D | 0.79 | 0.81 | 0.74 | 0.77 |
| Set E | 0.86 | 0.85 | 0.83 | 0.85 |
| Set F | 0.85 | 0.85 | 0.83 | 0.84 |
| Set G | 0.85 | 0.87 | 0.84 | 0.85 |
| Set H | 0.81 | 0.83 | 0.83 | 0.82 |
| Set I | 0.87 | 0.86 | 0.84 | 0.86 |
| Set J | 0.75 | 0.81 | 0.79 | 0.78 |

**Table 7.**
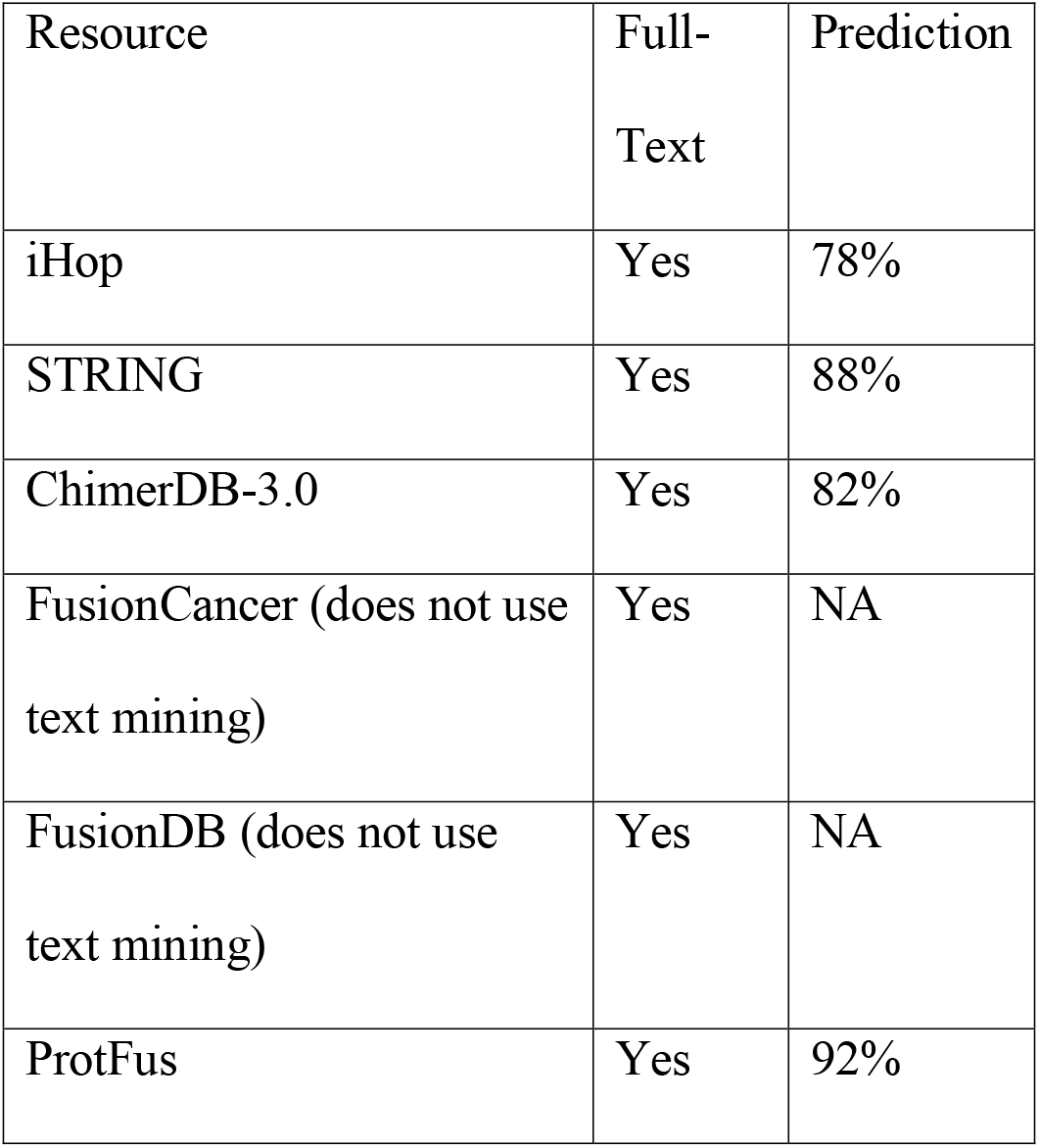
Performance of ProtFus in comparison to other resource.

| Resource | Full-Text | Prediction |
| --- | --- | --- |
| iHop | Yes | 78% |
| STRING | Yes | 88% |
| ChimerDB-3.0 | Yes | 82% |
| FusionCancer (does not use text mining) | Yes | NA |
| FusionDB (does not use text mining) | Yes | NA |
| ProtFus | Yes | 92% |

### 3.3 Entity recognition from fusion and PPI corpus

Moreover, the back-end ‘Synonyms’ (fusion corpus) consists of #entity# #relation token# like #fusion# #fusions, fusion transcript, fusion transcripts, fusion protein, fusion proteins, fusion gene, fusion genes#, whereas ‘Synonyms’ (ppi corpus) consists of #entity# #relation token# like #Activate# #activate, activates, activiated, activating, activation, activator#. Similarly, the back-end ‘Synonyms’ (fusion) consists of #Fusion proteins# #Synonyms# #Alternate representations# like #EWS-FLI1# #SH2D1B EAT2, ELK1, PDGFC SCDGF, ETV2 ER71, ETSRP71# #ews fli1, EWSR1 EWS, EWSR1/FLI1, EWS FLI-1#. The ‘parser’ and entity recognition module uses ‘Rulebase’ and ‘Short Form Recognition’ back-end resources for identifying the final ‘best suited’ entities and tokens as well as filtering out the false positives. The ‘Rulebase’ (for normalization) consists of #Rule# #Input# #Output# #Reg Ex# like #Normalization of case# #BCR-ABL, bcr-abl, BCR:ABL, bcr:ABL, BCR/ABL, bcr/abl# #BCR-ABL, BCR:ABL, BCR/ABL#. Similarly, the ‘Rulebase’ (for regular expression) consists of #Characteristics# #Description# #Rule# #Reg Ex# like #Fusion token# #Tokens with fusion word occurrence# #Should be separated by space/tokens# #(‘fusion|fusions|fusion genes|gene fusion|fusion protein|fusion transcripts’)# etc. Taken together, this process indicate that ProtFus considers all possible combinations of representing the fusions in text, by considering the back-end Rulebase as well as the fusion and ppi corpus.

To date, a limited number of tools exist which work on extracting and identifying fusion proteins as well as their interactions from literature. We tested ProtFus on identifying mentioned occurrences of 358 fusion proteins (based on Mitelman Database) from PubMed articles, out of which we provide the result statistics of top 100 fusion proteins based on their identification from text (***Table S5, Suppl. data***). For example, in case of BCR-ABL1 fusion protein (PubMed ID = 9747873), ProtFus identifies its occurrence as ‘Both Bcr-Abl fusion proteins exhibit an increased tyrosine kinase activity and their oncogenic potential has been demonstrated using in vitro cell culture systems as well as in in vivo mouse models’ (***Table S5, Suppl. data***). Similarly, ProtFus also identifies interactions among fusion proteins (***Table S6, Suppl. data***), such as in case of BCR-ABL1 fusion protein (PubMed ID = 9747873), ‘The SH2-containing adapter protein GRB10 interacts with BCR-ABL’ (***Table S6, Suppl. data***). The essential parameters to check the accuracy of text-mining based algorithms is to identify Precision, Recall and F-Score. ProtFus identifies fusion proteins with Precision ranging from 0.33 to 1.0 (average=0.83), Recall = 0.4 to 1.0 (average=0.84) and F-Score = 0.4 to 1.0 (average=0.81), whereas for protein-protein interactions, Precision = 0.42 to 1.0 (average=0.81), Recall = 0.5 to 1.0 (average=0.81) and F-Score = 0.59 to 1.0 (average=0.83) respectively. Thus, the overall accuracy of ProtFus allows it to extract the functional as well as key attributes for fusions and their interaction appearances in the text.

### 3.4 Training and Testing

Now, we used a classical Naïve Bayes algorithm for performing training as well as prediction. The datasets were partitioned based on known fusions and their interactors from literature that acted as training (40%) (***Table 1***). The rest of the data (around 60%) were used for testing from all PubMed references (2013-2017) (***Table 2***). There was no overlap among training and testing data. The screening was done based on distinct PubMed IDs. This is followed by modeling the decisions for assigning labels to raw input data. This kind of classification algorithms can also be thought of a convex optimization problem, where one needs to identify the minima of a convex function *ρ* associated with an input vector *v*, having *n* entries Eq (2),

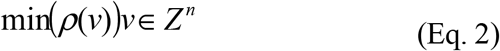

Here, the objective function can be defined as Eq (3),

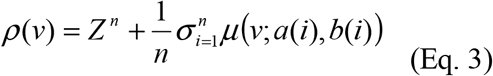

Here, vectors *a*(*i*) ∈ *Z*^n^ are training instances (1 ≤≤ *n*), *y*(*i*) ∈ *Z*^n^ act as labels. For checking the accuracy of our algorithm, we consider a five 10-fold cross-validation. For this purpose, we partition the input text into ten equal-sized sub-samples, of which five were retained as testing while five were used for model building. We also used the standard Precision, Recall and F-score values for validating the results. Precision ( *P*) is defined as the fraction of retrieved instances that are relevant to the study. It can also be defined as the probability that a randomly selected retrieved information is relevant Eq (4).
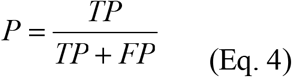

Here, *TP* = true positive and *FP* = false positive. Similarly, Recall ( *R*) is defined as the fraction of relevant instances that are retrieved for the study. It can also be defined as the fraction of the information that are relevant to the query that are successfully retrieved Eq (5).
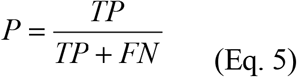

Here, *FN* = false negative. Finally, *F* – *score* is the harmonic mean of precision and recall Eq (6).
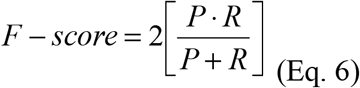

For example, if the standard query text contains 3 tokens that could be categorized as fusions, and ProtFus identifies 2 out of it, the accuracy can be calculated as: True (standard) tokens = *n*, *y*, *n*, *a*; Predicted (by ProtFus) tokens = *n*, *n*, *n*, *a* (here, *n* = no token instance, *y* = token instance, *a* = noise). In this case, *Precision* = 0:75, *Recall* = 0:75, *F-score* = 0:75, respectively. Similarly, the corresponding accuracy plot can also be drawn by providing information about *Precision*, *Recall*, *F-score* values and the number of runs. Thus, ProtFus provides this facility for users to visually consider the accuracy of their searches.

### 3.5 Big Data processing using ProtFus and ChiPPI

Mining biomedical texts generates a considerable amount raw as well as informative data, where the key challenge is managing as well as storing them for data analysis tasks. For this purpose, we built the Protein-Protein Interaction of Fusions (PPI-Fus) database (ppi-fus.md/biu.ac.il/bin/protfusdb.pl), supported by Apache Tomcat and My-SQL. It is an open source Big Data processing framework that supports ETL (Extract, Transform and Load), machine learning, as well as graph generation. Finally, some classical text mining tasks can also be performed by identifying biological, functional, literals, and miscellaneous tokens, as well as chunks from text. The word-token tagger has a back-end Synonyms (with synonyms resource) whereas the RegEx tagger has a back-end Synonyms (with rulebase). This is followed by parsing the tokens for performing entity recognition. In the interface, the search can be using PubMed or uploading a text file or based on a specific input text. For example, in case of an input text, the result is displayed in a separate window, with the fusion proteins being highlighted.

Further, in case of identifying protein-protein interactions among fusions, the result window includes the input text with interactions being highlighted. Another feature of ProtFus includes directly searching using PubMed articles. Users can select from the given drop-down box, the number of articles that need to be considered for searching fusions and their interactions. The result includes the abstracts of all those articles which match best with fusion proteins keywords. This file can be further used for highlighting the fusions and their interactions. These results indicate that our text mining method successfully identifies unique novel fusion protein and their interactions from text by tagging tokens, that act as entities.

Considering discrete protein domains as binding sites for specific domains of interacting proteins, we have catalogued the protein interaction networks for more than 11,000 cancer fusions in order to build the Chimeric Protein-Protein-Interactions (ChiPPI) [35]. Mapping the influence of fusion proteins on cell metabolism and protein interaction networks reveals that chimeric protein-protein interaction (PPI) networks often lose tumor suppressor proteins, and gain onco-proteins. As a case study, we compared the results generated by ProtFus with the interaction prediction accuracy of ChiPPI [35]. For example, in BCR-JAK2 fusion, ProtFus provides multiple hits regarding its occurrence in literature, such as, “It was demonstrated preclinical studies that BCR-JAK2 induces STAT5 activation elicits BCRxL gene expression” (PMC3728137), as correctly predicted by ChiPPI (***Fig 4***).

### 3.6 ProtFus performs much better in compared to other text-mining resources

***Table 7*** represents the accuracy of ProtFus as compared to iHop [10, 3], STRING [33], ChimerDB-3.0 [18], FusionCancer [22] and FusionDB [34] resources. The iHOP method is based on a dictionary approach, wherein abstracts are searched for gene synonyms using hashcodes, followed by assigning genes to precise text positions. STRING parses large scientific texts from various resources like SGD, OMIM, FlyBase, and PubMed to search for statistically relevant co-occurrences of gene names. ChimerDB-3.0 chooses fusion gene candidate sentences from PubMed which are further used for training a machine learning model. FusionCancer and FusionDB do not use text mining for fusion prediction, but we used them as a resource-based comparison for predicted fusions. The efficiency of our algorithm is around 92%. We also found the Receiver Operating Characteristic (ROC) curves [29] for quantitative representation of our method. ***Fig 3*** shows representative ROC curves generated in a typical experiment using ‘abstracts’ data. In compared to full-text articles, the prediction was better for abstracts. This is due to the fact that the size of feature space is too large for full-text articles. For text classification purposes, abstracts may work better than full-text scientific articles.

**Fig 3:**
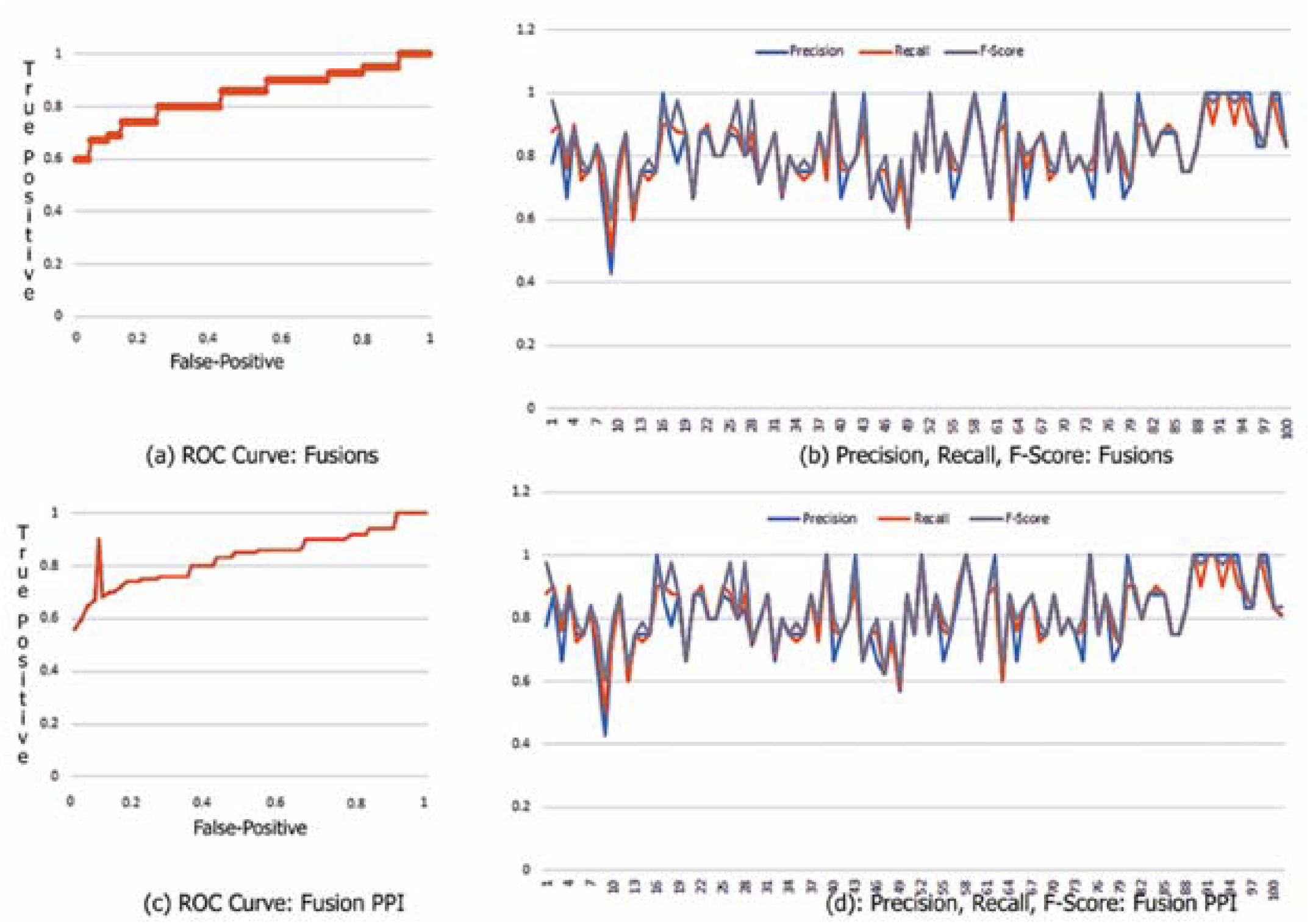
ROC curve for Naïve Bayes and Accuracy. The ROC curves for Naïve Bayes; Fusion and Fusion PPI detection; Precision, Recall and F-Score rates.

**Fig 4:**
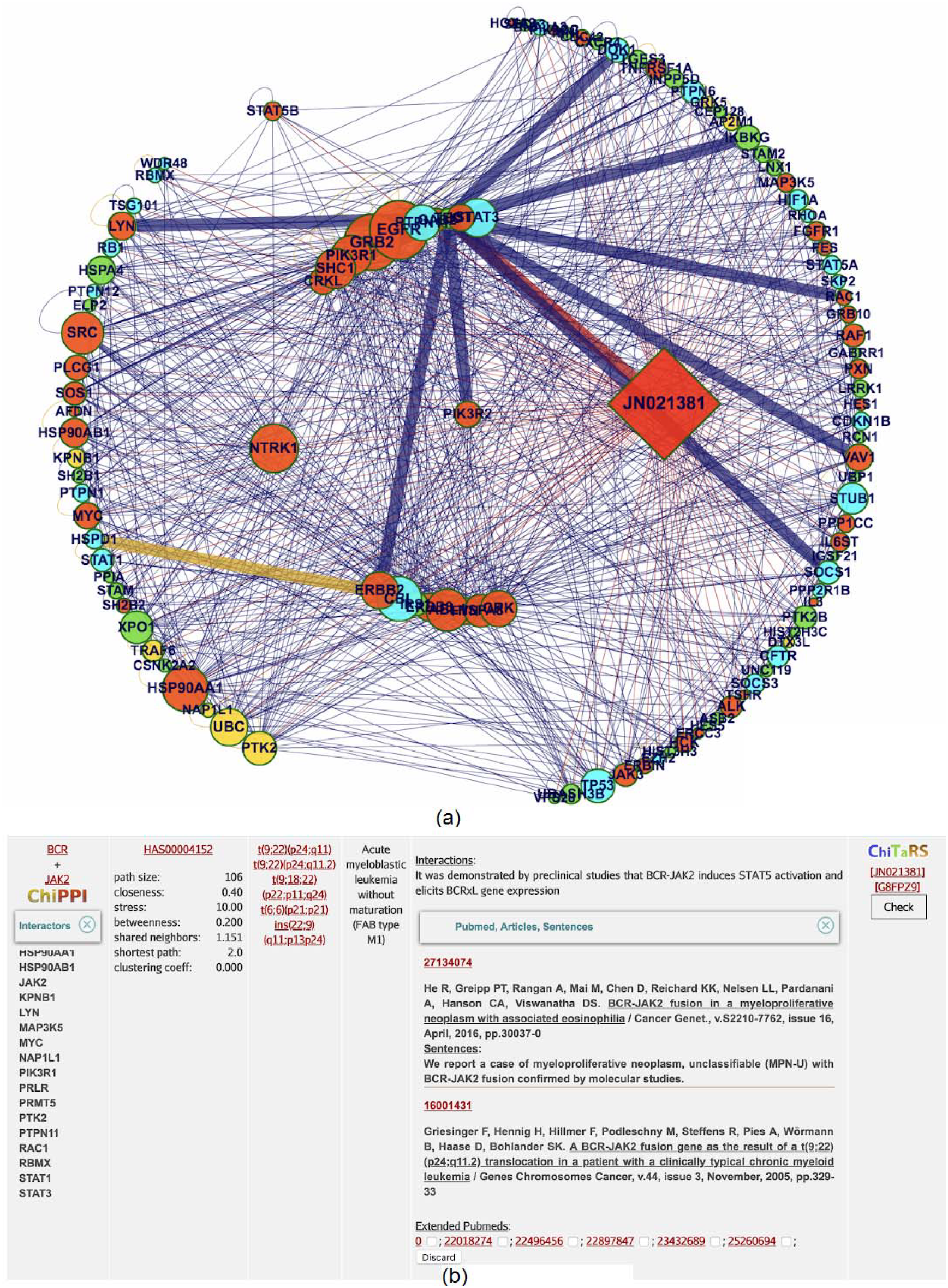
ChiPPI analysis (a) PPI-Fus/ProtFus prediction for BCR-JAK2 and STAT5B interaction (b) as predicted by ProtFus.

## Discussions and Conclusions

This study focused on investigating large-scale biomedical text classification downloaded from PubMed. We utilized classical text-mining, machine learning strategies, as well as Big Data infrastructure to design and develop a distributed and scalable framework. This was used to extract identify fusion proteins and their interactions for classifying information extracted from tens of thousands of abstracts and/or full-text articles associated MeSH terms. The accuracy of predicting a cancer type by Naïve Bayes using the abstracts was 92%, while its accuracy using the 103,908 abstracts (for fusions only); 90,639 full texts (for fusions only); 185,606 abstracts (for fusion protein interactions); 353,535 full texts (for fusion protein interactions) was 88%. This study demonstrates the potential for text mining of the large-scale scientific articles on a novel Big Data infrastructure, with the real-time update from new articles published daily. Therefore, ProtFus can be extended to other areas of biomedical research for example, the patients’ drug response, in order to improve the medical data mining in the Personalized Medical approaches.

## Acknowledgements

We thank Dr. Dorith Raviv-Shay for proofreading the manuscript. Prof. Alfonso Valencia, Dr. Michael Blank and Dr. Hava Gill-Henn for the viable discussions on the topic.

## Supporting Information

Table S1: Root and relation tokens, Bibliography

Table S2: Action Tokens for Fusion PPI

Table S3: Synonyms for Fusions, Dictionary

Table S4: Rulebase

Table S5: Fusion tokens identified by ProtFus for 100 PubMed IDs

Table S6: Fusion PPIs tokens identified by ProtFus for 100 PubMed IDs

